# The structural basis for cancer drug interactions with the catalytic and allosteric sites of SAMHD1

**DOI:** 10.1101/296624

**Authors:** Kirsten M. Knecht, Olga Buzovetsky, Constanze Schneider, Dominique Thomas, Vishok Srikanth, Florentina Tofoleanu, Krystle Reiss, Nerea Ferreirós, Gerd Geisslinger, Victor S. Batista, Xiaoyun Ji, Jindrich Cinatl, Oliver T. Keppler, Yong Xiong

## Abstract

SAMHD1 is a deoxynucleoside triphosphate triphosphohydrolase (dNTPase) that depletes cellular dNTPs in non-cycling cells to promote genome stability and to inhibit retroviral and herpes viral replication. In addition to being substrates, cellular nucleotides also allosterically regulate SAMHD1 activity. Recently, it was shown that high expression levels of SAMHD1 are also correlated with significantly worse patient responses to nucleotide analogue drugs important for treating a variety of cancers, including Acute Myeloid Leukemia (AML). In this study, we used biochemical, structural, and cellular methods to examine the interactions of various cancer drugs with SAMHD1. We found that both the catalytic and the allosteric sites of SAMHD1 are sensitive to sugar modifications of the nucleotide analogs, with the allosteric site being significantly more restrictive. We crystallized cladribine-TP, clofarabine-TP, fludarabine-TP, vidarabine-TP, cytarabine-TP, and gemcitabine-TP in the catalytic pocket of SAMHD1. We find that all of these drugs are substrates of SAMHD1 and that the efficacy of most of these drugs is affected by SAMHD1 activity. Of the nucleotide analogues tested, only cladribine-TP with a deoxyribose sugar efficiently induced the catalytically active SAMHD1 tetramer. Together, these results establish a detailed framework for understanding the substrate specificity and allosteric activation of SAMHD1 with regards to nucleotide analogues, which can be used to improve current cancer and antiviral therapies.

**Significance:** Nucleoside analogue drugs are widely used to treat a variety of cancers and viral infections. With an essential role in regulating the nucleotide pool in the cell by degrading cellular nucleotides, SAMHD1 has the potential to decrease the cellular concentration of frequently prescribed nucleotide analogues and thereby decrease their clinical efficacy in cancer therapy. To improve future nucleotide analogue treatments, it is important to understand SAMHD1 interactions with these drugs. Our work thoroughly examines the extent to which nucleotide analogues interact with the catalytic and allosteric sites of SAMHD1. This work contributes to the assessment of SAMHD1 as a potential therapeutic target for cancer therapy and the future design of SAMHD1 modulators that might improve the efficacy of existing therapies.

## Introduction

The sterile alpha motif and histidine-aspartate domain–containing protein 1 (SAMHD1) is a phosphohydrolase that severs the triphosphate group from deoxynucleoside triphosphates (dNTPs) (1, 2). A major function of SAMHD1 is to reduce the dNTP pool in non-cycling cells, making it an important regulator of dNTP levels in the cell (1). In addition to its role in regulating genome stability, SAMHD1 is best known for its ability to block infection of a broad range of retroviruses including human immunodeficiency virus type 1 (HIV-1). During viral infection, SAMHD1 depletes the cellular dNTPs needed for reverse transcription of the viral RNA genome (2-8). In the cell, SAMHD1 activity is modulated by allosteric activation and phosphorylation (9-19). Nucleotide binding to two allosteric sites of each subunit leads to the assembly of activated SAMHD1 tetramer. Allosteric site (Allo-site) 1 only accommodates guanosine bases (GTP or dGTP), but any canonical dNTP can bind Allo-site 2 (15, 18, 20-22). When dNTPs are needed for DNA synthesis, phosphorylation at residue T592 destabilizes the active tetramer of SAMHD1 thereby downregulating SAMHD1 activity (9, 10, 13).

SAMHD1 is an important general sensor and regulator of the dNTP pools and thus it is crucial to genome maintenance. All canonical dNTPs are both substrates and allosteric activators, and this promiscuity allows SAMHD1 to target therapeutic molecules that resemble nucleotides in structure. Nucleoside analogues are a large class of drugs which are used to treat viral infections and many types of cancers (23-30). These compounds interfere with viral replication or cancer cell proliferation upon incorporation into newly synthesized DNA, resulting in chain terminations, accumulation of mutations, and often cell apoptosis (24, 27, 28). Several recent reports have demonstrated that SAMHD1 reduces the efficacy of nucleotide analogue drugs by depleting their cellular concentrations (31, 32). Strikingly, SAMHD1 expression levels were shown to be highly predictive of patient response to cytarabine, the primary treatment for acute myeloid leukemia (AML) (31). Better characterization of SAMHD1 substrate specificity is needed for the optimal administration of current nucleoside analogue drugs and the development of new compounds for antiviral and cancer therapies that resist SAMHD1 degradation.

Interestingly, SAMHD1 activity has been shown to increase the efficacy of some nucleotide analogues that are not substrates of SAMHD1 (33, 34). In these cases, SAMHD1 activity allows the non-substrate nucleotide analogues to better compete for target active sites by depleting cellular dNTPs. Therefore, depending on how SAMHD1 interacts with a particular nucleotide analogue of interest, it might be desirable to either inhibit or increase SAMHD1 activity. To selectively modulate SAMHD1 activity, it is important to fully understand how nucleotide analogue drugs either bind to the allosteric site to assemble an activate tetramer or bind the catalytic site to be hydrolyzed.

In this study, we characterized SAMHD1 interactions with a panel of nucleotide analogues that are used to treat a variety of cancers and viral infections (23, 26, 27, 29, 35, 36) (Fig. 1A). We found that the catalytic site of SAMHD1 is very promiscuous, allowing SAMHD1 to hydrolyze most of the analogues tested here. On the other hand, Allo-site 2 is more restrictive to modifications of the 2’ sugar moiety of the drug. These results are important for the assessment of SAMHD1 as a potential therapeutic target for cancer therapy, the design of non-hydrolysable derivatives, and the development of modulators of SAMHD1 activity to combine with existing therapies. In addition, this work contributes to a greater understanding of the structural and biochemical principles of SAMHD1 substrate selectivity and allosteric activation.

**Figure 1.**
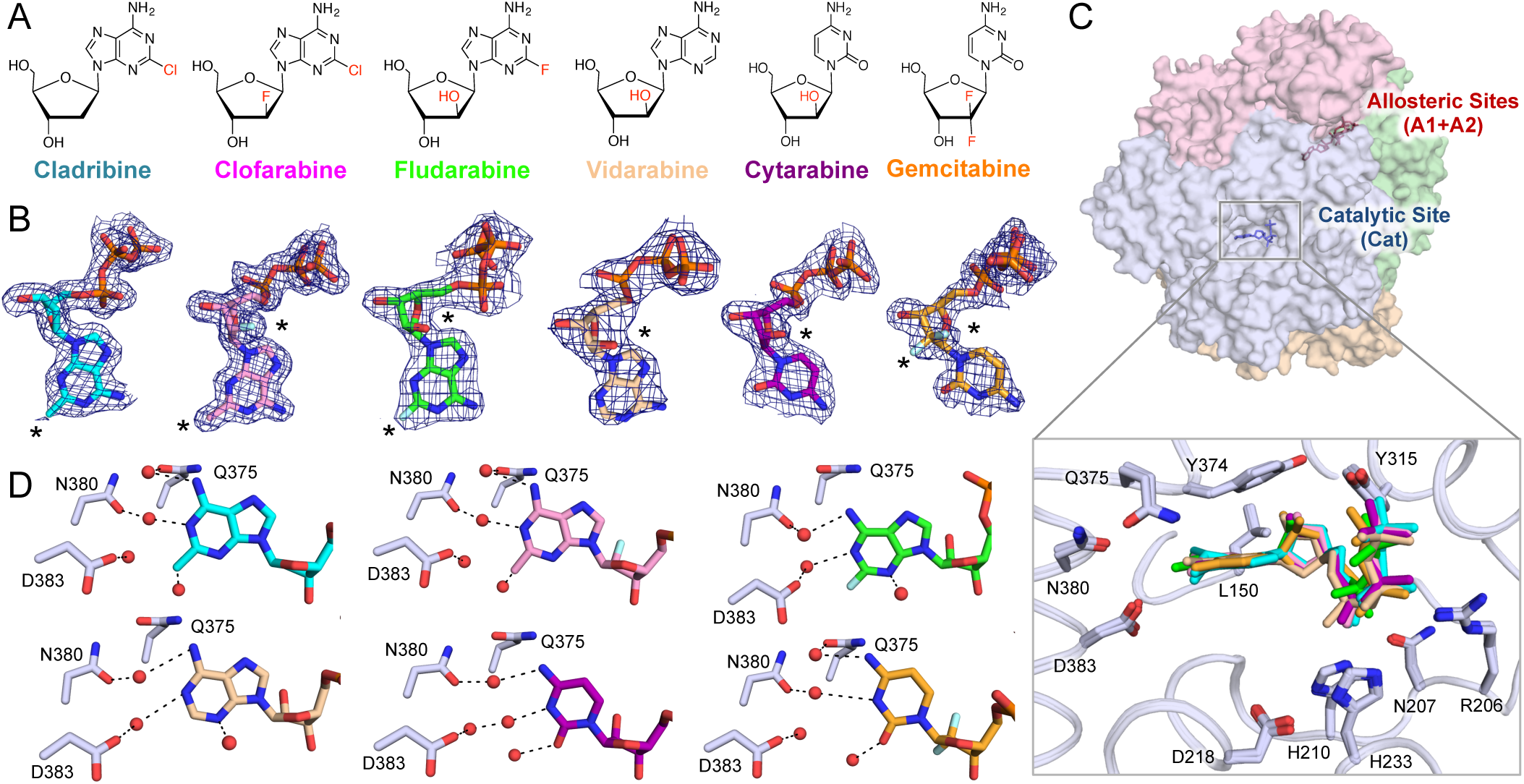
Substrate specificity of SAMHD1 is determined by 2’ sugar moiety. (A) Chemical structures of nucleoside analogues used in this study. (B) 2F_o_-F_c_ electron density (σ = 1.0) for the nucleotide analogue drugs crystallized in the catalytic pocket of SAMHD1. Black asterisks indicate sites of modifications. (C) Top: Transparent surface view of SAMHD1 tetramer with each subunit in a different color. Selected allosteric nucleotides are shown in red sticks and a nucleotide in a catalytic pocket is shown in blue sticks. Bottom: Superposition of all the nucleotide analogues bound to the SAMHD1 catalytic pocket. SAMHD1 backbone is shown as coils with side chains shown as sticks. Cladribine-TP (cyan), clofarabine-TP (magenta), fludarabine-TP (green), vidarabine-TP (wheat), cytarabine-TP (purple), and gemcitabine-TP (orange) are shown as sticks. (D) Water networks (shown as red spheres) observed for each nucleotide analogue bound to the SAMHD1 catalytic site. Black dashed lines indicate hydrogen bonds.

## Results

### Crystal structures of nucleotide analogues bound to the SAMHD1 catalytic pocket

To better understand the structural basis for nucleotide analogue binding in the catalytic pocket of SAMHD1, we co-crystallized the inactive catalytic domain of SAMHD1 (residues 113-626 with H206R/D207N mutations to inhibit catalysis) with six selected cancer and antiviral drugs. This SAMHD1 mutant has been previously shown to be identical in conformation and nucleotide-binding properties to the WT enzyme, but it is more amenable to crystallization (18, 19, 21, 22). The crystal structures of these SAMHD1-nucleotide analogue complexes were determined at resolutions ranging from 1.7 Å to 2.5 Å (Table 1), with electron density that allows for unambiguous identification of each substrate in the catalytic pocket (Fig. 1B). The structures of SAMHD1 bound to these nucleotide analogues are similar to those obtained with canonical nucleotides (19, 21). Nucleotide analogue binding does not alter the architecture of the SAMHD1 catalytic pocket (Fig. 1C), and these nucleotide analogues adopt similar conformations to the canonical nucleotides in the pocket. This suggests that the nucleotide modifications tested here do not disturb the overall integrity of the catalytic pocket or induce large structural rearrangements. The promiscuous catalytic pocket of SAMHD1 likely accommodates other nucleotide analogues with similar modifications.

**Table 1.**
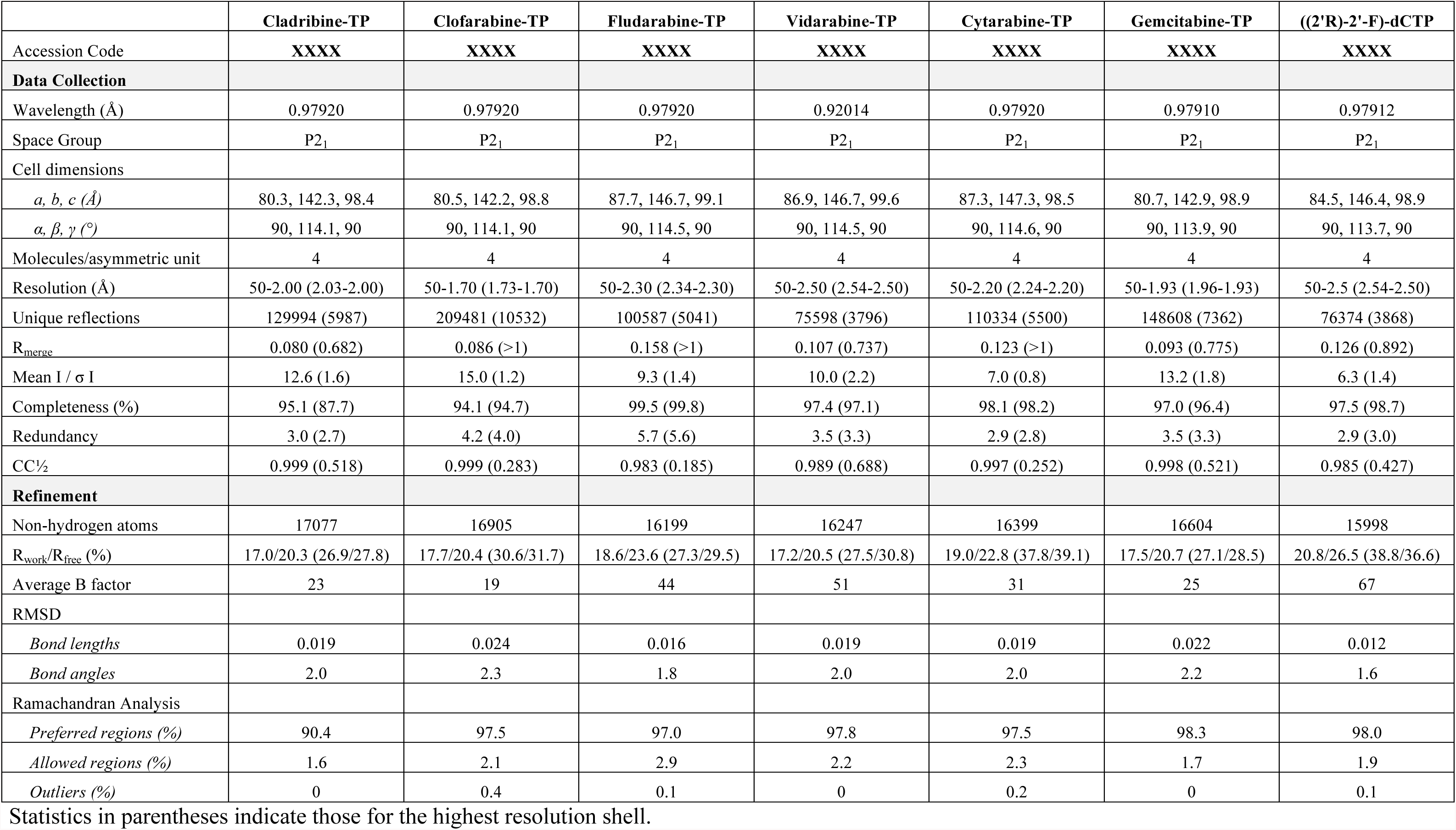
Data collection and refinement statistics for the seven crystal structures of SAMHD1 HD bound to nucleotide analogues.

### Base modification has modest effects on substrate binding

Modifications on the Watson-Crick base edge are well tolerated at the catalytic site of SAMHD1. Consistent with previous observations with dNTPs (19, 21), there are no base-specific interactions between SAMHD1 and the bases of the nucleotide analogues. Water networks in the active site of SAMHD1 stabilize the nucleotide analogues in the pocket, as for canonical dNTPs. In each case, three to four water molecules bridge the interaction between SAMHD1 and the Watson-Crick and sugar edges of the bound analogue (Fig. 1D). Cytarabine-TP and gemcitabine-TP, each containing an unmodified cytosine base, maintain water network interactions that closely resemble those reported for dCTP (Fig. 1D) (19). Surprisingly, we found that the dATP derivatives (cladribine-TP, clofarabine-TP, fludarabine-TP, and vidarabine-TP) seem to bind the catalytic pocket more stably than dATP. In our previous structural studies, the adenosine base was not resolved in the dATP-SAMHD1 co-crystal structure due to its relatively weak binding affinity to the catalytic site compared to other dNTPs (19). However, clear electron density for the bases of clofarabine-TP, cladribine-TP, fludarabine-TP, and vidarabine-TP were observed in the current structures, suggesting that these analogues have higher binding affinity for the active site than that of dATP (Fig. 1B). The additional chlorine atom (cladribine-TP and clofarabine-TP) or fluorine atom (fludarabine-TP) at the C2 position of the bases contribute to a more extensive water network and stabilize these molecules in the active site (Fig. 1D). These results suggest that small modifications in the base are not likely to disrupt binding to the SAMHD1 catalytic pocket because water networks that contact the base are flexible. Some nucleotide modifications may even enhance the water network surrounding the nucleotide analogue and possibly increase its binding affinity.

### 2’ sugar moiety regulates SAMHD1 substrate specificity

It is well established that NTPs with ribose sugars ((2’R)-2’-OH) are not substrates of SAMHD1 (2, 19). We examined how sugar modifications found in nucleotide analogue drugs influence their binding to and hydrolysis by SAMHD1, as it has been shown that arabinose-based nucleotides with 2’S sugar modifications are substrates of SAMHD1 (31, 32). Our crystal structures corroborate the finding that arabinose-based sugars, with either ((2’S)-2’-OH) or ((2’S)-2’-F) modifications, are allowed in the catalytic site of SAMHD1. In addition to the canonical interactions observed with dNTP substrates (19), we observe van der Waals and stacking interactions between active site residues Y315 and Y374 and the ((2’S)-2’-OH/F) atoms of cytarabine-TP, clofarabine-TP, fludarabine-TP, vidarabine-TP and gemcitabine-TP in the catalytic pocket (Fig. 1C, 2A and 2B). Since the arabinose-like sugar modifications provide an additional interaction with SAMHD1, these modifications likely stabilize these analogues in the catalytic pocket. In contrast, we crystallized SAMHD1 in the presence of 10 mM ((2’R)-2’-F)-dCTP, which has a ribose-like modification, but electron density for this nucleotide analogue was not observed in the catalytic pocket (Fig. S1A).

Our activity assays also confirmed that the substrate specificity of SAMHD1 has a general dependency on the stereochemistry of the sugar moiety at the 2’ position, allowing the arabinose-like 2’S, but not the ribose-like 2’R geometry to be hydrolyzed. We used a malachite green assay to measure the rate of hydrolysis of each drug by pre-assembled SAMHD1 tetramers (37). Consistent with our crystallographic data, all of the drugs we observed in the SAMHD1 co-crystal structures were hydrolyzed by SAMHD1 at various rates, whereas the non-binding ((2’R)-2’-F)-dCTP was not hydrolyzed (Fig. S1B). Overall, the dATP analogues were hydrolyzed at higher rates than those of the dCTP analogues, consistent with SAMHD1’s known preference for dATP as a substrate over dCTP (19). Although some 2’ sugar modifications are tolerated, modifications larger in size were associated with slower rates of hydrolysis. Cladribine-TP with an unmodified deoxyribose sugar was hydrolyzed by SAMHD1 at the highest rate, while gemcitabine-TP with a 2’,2’-difluorine sugar modification was hydrolyzed at a lower rate (Fig. S1B). These results confirmed other reports that arabinose-based nucleotide analogues are substrates of SAMHD1, whereas the ribose-based ((2’R)-2’-F)-dCTP is not (31, 32).

Surprisingly, gemcitabine-TP was hydrolyzed by SAMHD1, albeit at a slow rate (Fig. S1B). Gemcitabine-TP had previously been reported not to be a substrate of SAMHD1 (32). Hollenbaugh *et al.* showed that the concentration of gemcitabine-TP in monocyte-derived macrophages was not dependent on SAMHD1 protein levels (32). This is perhaps due to the difficulty in detecting the low hydrolysis rate in the complex cellular environment. To reconcile this discrepancy, we directly compared the hydrolysis of the doubly modified gemcitabine-TP to the singly modified ((2’S)-2’-F)-dCTP and ((2’R)-2’-F)-dCTP *in vitro* in a time course assay (Fig. 2C). We observed that SAMHD1 indeed hydrolyzes gemcitabine-TP at an intermediate rate between ((2’S)-2’-F)-dCTP and the unreactive ((2’R)-2’-F)-dCTP (Fig. 2C). As evidenced by the electron density of gemcitabine-TP in the catalytic pocket, both the 2’S and 2’R fluorine modifications are accommodated in the catalytic pocket of SAMHD1 (Fig. S1C). However, the substructure ((2’R)-2’-F)-dCTP, with a single 2’R modification, is not accommodated. Modeling of this compound into the catalytic pocket predicts that the (2’R)-2’-F atom would clash with L150 (Fig. 2D). This suggests that the 2’R modification is permitted only in the context of a 2’,2’-difluorine sugar modification, but not alone (Fig. 2B and 2D). SAMHD1’s discrimination between these two substrates may arise from interactions between Y315/Y374 and the 2’S fluorine atom, which might partially compensate for the sterically unfavorable 2’R fluorine modification (Fig. 2B and 2D). Although gemcitabine-TP is allowed in the catalytic pocket, the presence of the 2R’ modification may lead to a less suitable positioning of the nucleotide in the catalytic pocket for catalysis as compared to 2’S modifications alone (Fig. 2A and Fig. 2B). These results support the notion that the catalytic pocket of SAMHD1 is highly sensitive to the stereochemistry of 2’ sugar modification.

**Figure 2.**
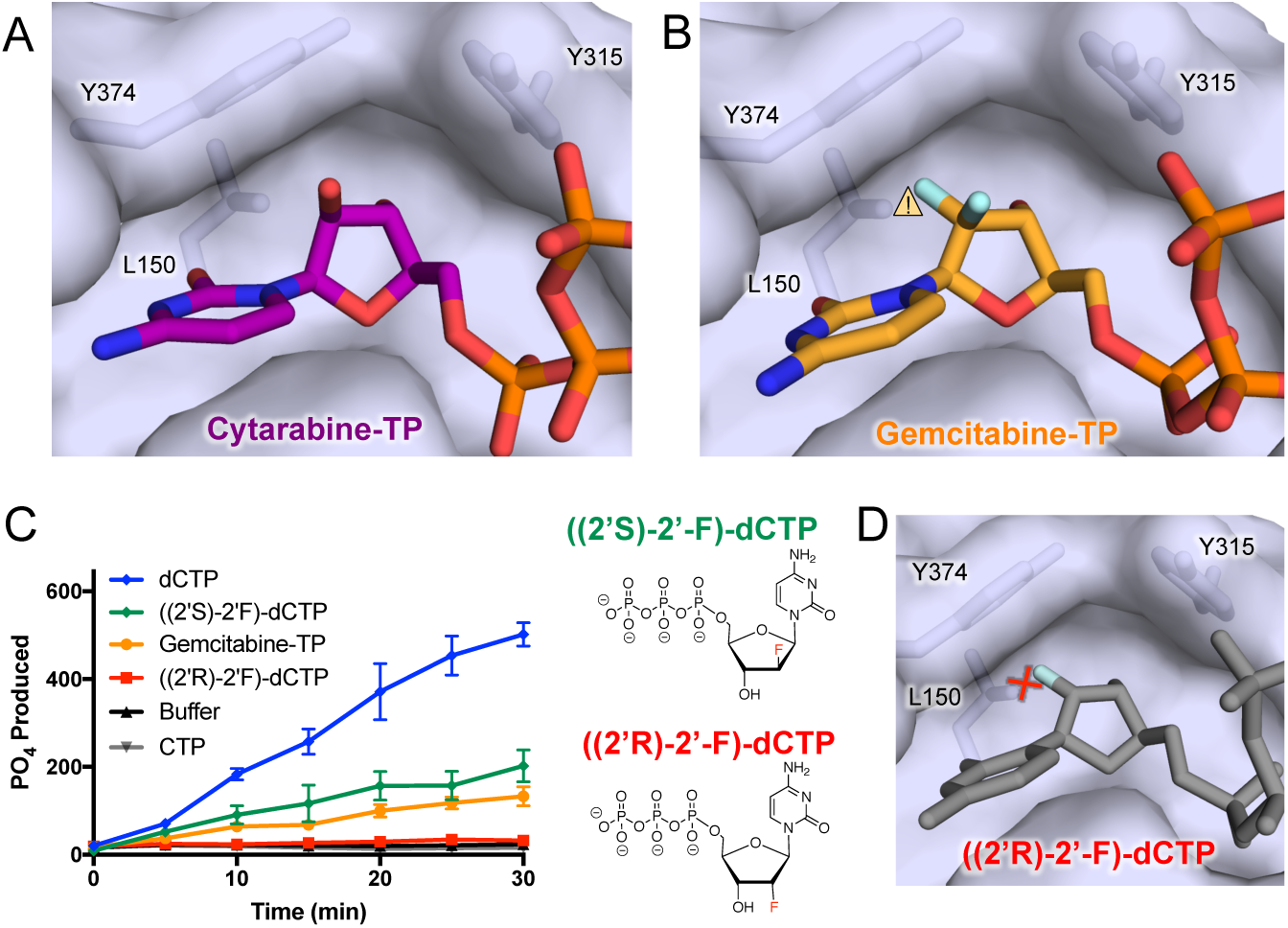
Gemcitabine-TP but not ((2’R)-2’-F)-dCTP is hydrolyzed by SAMHD1 *in vitro*. (A) ((2’S)-2’-OH)) of cytarabine-TP is stabilized by residues Y374 and Y315 through van der Waals interactions. Transparent surface of SAMHD1 is shown with key residues in sticks. (B) 2’,2’-difluorine sugar modification of gemcitabine-TP is stabilized by Van der Waals interactions with residues Y374 and Y315 to compensate potential close contact (yellow caution triangle) between ((2’R)-2’-F) atom and residue L150 in the catalytic site. (C) Left: dNTPase activity of SAMHD1 over the course of 30 minutes was measured using a malachite green assay. Product is normalized to SAMHD1 concentration (nmol PO_4_ / nmol SAMHD1). SAMHD1 tetramers were pre-assembled with 250 uM GTP and dATP and then diluted 100-fold into 125 uM gemcitabine-TP, dCTP, CTP, ((2’S)-2’-F)-dCTP, ((2’R)-2’-F)-dCTP, or buffer. Error bars represent standard error of the mean (SEM) of three independent experiments. Right: chemical structures of ((2’S)-2’-F)-dCTP and ((2’R)-2’-F)-dCTP analogues. (D) ((2’R)-2’-F)-dCTP (gray sticks) modeled into the catalytic pocket potentially clashes (red cross) with residue L150.

### SAMHD1 hydrolyzes various nucleotide analogues *in vivo*

Since SAMHD1 is capable of hydrolyzing cancer drugs *in vitro*, it has the potential for decreasing their therapeutic efficacies. We explored the extent to which the triphosphates of each of these nucleotide analogues are degraded by SAMHD1 in the cell following uptake and metabolic activation. A recent report demonstrated a strong inverse correlation between SAMHD1 expression in leukemic blasts and AML patients’ clinical response to cytarabine therapy (31). Similarly, we found that the IC_50_ values of fludarabine-TP and clofarabine-TP in AML cell lines were also correlated with SAMHD1 protein expression levels (Fig. 3A). However, this effect was not observed for cladribine-TP or gemcitabine-TP. We also tested whether the depletion of SAMHD1 in THP-1 cells via knock-out or targeted proteasomal degradation by Vpx-VLPs affected the IC_50_ values of these drugs. Although a strong effect was observed for cytarabine-TP and a moderate effect was observed for fludarabine-TP, clofarabine-TP, vidarabine-TP and cladribine-TP (Fig. 3 and Fig. S2A), there was no effect on the IC_50_ value for gemcitabine-TP (Fig. 3B). To directly measure SAMHD1’s effect on each nucleotide analogue’s concentration in the cell, we used LC-MS/MS to quantify drugs found in cells with or without SAMHD1 expression. Our results show that clofarabine-TP, fludarabine-TP, and cytarabine-TP were strongly depleted by SAMHD1, whereas cladribine-TP was depleted to a low extent and gemcitabine-TP levels remained unaffected (Fig. 3C). Although SAMHD1 hydrolyzes gemcitabine *in vitro* with a low activity, this effect may not be detectable under normal cellular conditions. Our findings corroborate with previous reports that gemcitabine-TP is not significantly degraded by SAMHD1 *in vivo* (31, 32).

**Figure 3.**
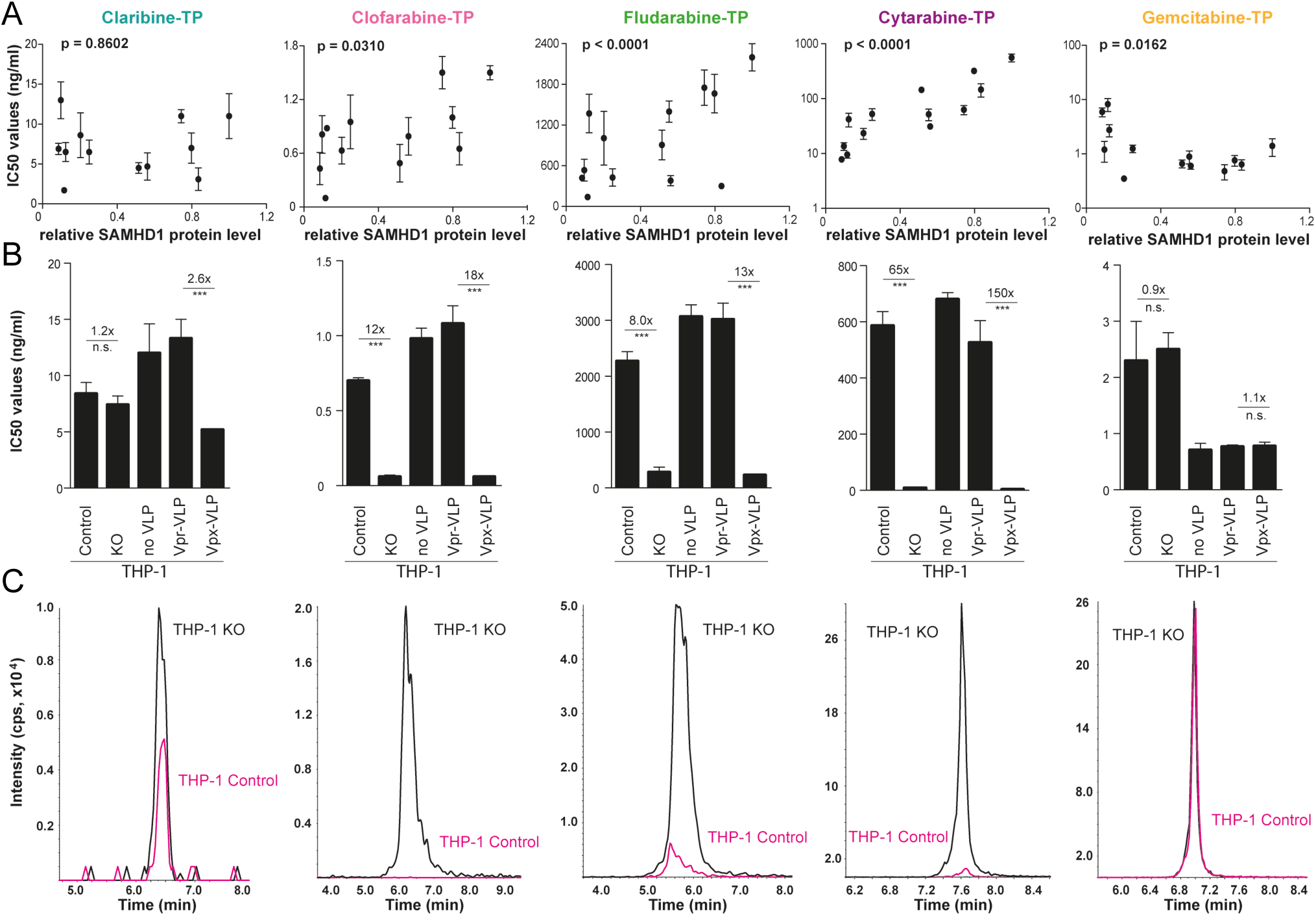
SAMHD1 depletes several TPs of nucleoside analogues *in vivo*. (A) Correlations of cytarabine, clofarabine, fludarabine, cladribine, and gemcitabine concentrations inhibiting 50% of cell viability (IC_50_) and relative protein expression levels of SAMHD1 in 13 AML cell lines. Relative expression levels (ratios of a SAMHD1/β-actin) are shown as arbitrary units (a.u.). Ratio of SAMHD1/β-actin for parental THP-1 cells is set to 1 and ratios of other cell lines are set relative to it. Closed circles represent mean values and error bars indicate SEM of three independent experiments. (B) Cytarabine, fludarabine, clofarabine, cladribine or gemcitabine IC_50_ values of THP-1 KO cells or THP-1 cells transduced with VLPs carrying either lentiviral Vpr (Vpr-VLPs, control) or Vpx proteins (Vpx-VLPs). The bars represent mean values and the error bars are SEM of three independent experiments. The numbers above indicate factor of decrease of the IC_50_ values in the absence of SAMHD1. (C) Representative liquid chromatography tandem mass spectrometry measurements (LC-MS/MS) of cytarabine-TP, fludarabine-TP, clofarabine-TP, cladribine-TP or gemcitabine-TP in THP-1 KO cells (black) and THP-1 control cells (red). Statistical analyses were performed using unpaired two-tailed Students’ t-test comparing treated samples with untreated control. ***p < 0.001.

The structural and biochemical framework established in this study also allows for the rational modeling of other known cancer drugs into the catalytic pocket of SAMHD1. For example, the IC_50_ of nelarabine-TP in AML cell lines relies on the expression level of SAMHD1, indicating that it is a substrate of SAMHD1 too (Fig. S2B). Modeling nelarabine-TP into the catalytic pocket of SAMHD1 predicts that it would fit into the catalytic pocket like any other arabinose-based nucleotide analogue, such as vidarabine-TP (Fig. S2C).

### Allosteric site 2 of SAMHD1 is more restrictive than its catalytic site

In addition to establishing SAMHD1’s substrate specificity for nucleotide analogues, we also tested whether these drugs were capable of binding to the allosteric sites to induce the catalytically active SAMHD1 tetramer. None of the nucleotide analogues tested here contained the guanosine base required for Allo-site 1 binding, thus they alone were not sufficient for SAMHD1 activation (Fig. S1D). To test which analogues bind Allo-site 2, we monitored the oligomerization state of SAMHD1 in the presence of GTP and each of the analogues. Previous studies indicated that clofarabine-TP is an activator of SAMHD1 (38), but cytarabine-TP is not (31, 32). Consistent with these reports, our size exclusion chromatography (SEC) assays showed that only clofarabine-TP and cladribine-TP caused a shift in the elution profile of SAMHD1 towards a higher molecular weight species (Fig. 4A), with cladribine-TP being more effective at inducing SAMHD1 oligomerization. To further investigate the effects of these drugs on the SAMHD1 oligomerization state, we used sedimentation velocity analytical ultracentrifugation (SV-AUC) as an independent measure of SAMHD1 oligomerization. As observed with the SEC assay, cladribine-TP efficiently assembled SAMHD1 tetramers. However, clofarabine-TP-induced SAMHD1 oligomerization was not detected by SV-AUC (Fig. 4B), likely due to experimental constraints requiring about 40-fold less nucleotide analogues compared to SEC. Furthermore, our activity assay confirmed that clofarabine-TP has some capacity to activate SAMHD1, but it is significantly less proficient than cladribine-TP (Fig. 4C). None of the other nucleotide analogues were able to activate SAMHD1. Together, these results suggest that only minor sugar modifications are permitted for SAMHD1 allosteric activators.

**Figure 4.**
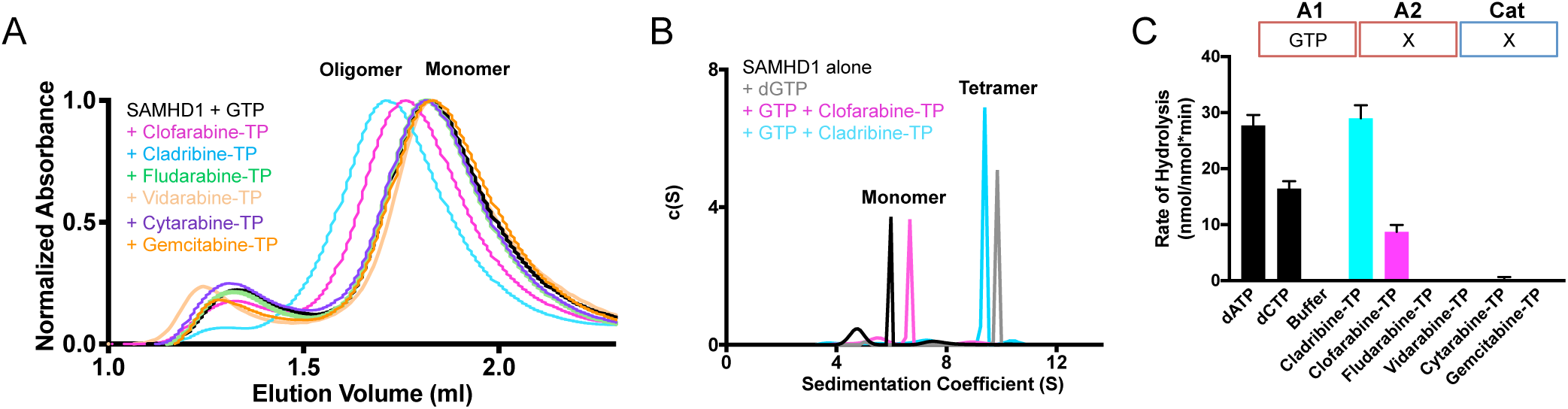
The allosteric sites of SAMHD1 are highly restrictive. (A) Size exclusion chromatography elution profile of SAMHD1 in the presence of 0.5 mM GTP and 4 mM of color-coded nucleotide analogue. (B) SV-AUC analysis of SAMHD1 in the absence of nucleotides or the presence of dGTP, GTP with clofarabine-TP, or GTP with cladribine-TP at a final concentration of 150 uM. (C) dNTPase assay performed in the presence of 125 uM GTP and 125 uM dCTP, dATP, nucleotide analogue, or buffer. Error bars represent SEM of three independent experiments.

### 2’ sugar modifications are highly restrictive at the allosteric site 2

To examine how SAMHD1 accommodates nucleotide analogues in the allosteric pocket, we attempted to crystalize SAMHD1 with GTP and each of the nucleotide analogues assayed above. Consistent with our activity assay and oligomerization measurements, only the two allosteric activators, cladribine-TP and clofarabine-TP, resulted in SAMHD1 tetramer crystals. The resulting structures revealed unambiguous electron density for each nucleotide in the Allo-site 2 (Fig. 5A). As predicted, the nucleotide analogues bind to Allo-site 2 and GTP binds to the adjacent Allo-site 1. The allosteric sites are not disturbed by cladribine-TP or clofarabine-TP, as the chlorine atom modification at the C2 position of the base does not interfere with the allosteric pocket interactions (Fig. 5B). Hydrogen bonds between the adenosine base and residues N119 and N358 (19) are preserved in the cladribine-TP and clofarabine-TP structures, allowing for the correct positioning of each nucleotide in the Allo-site 2 pocket. These results suggest that nucleotide analogues with some similar modifications at the Watson-Crick edge of the base may also be permitted in Allo-site 2.

**Figure 5.**
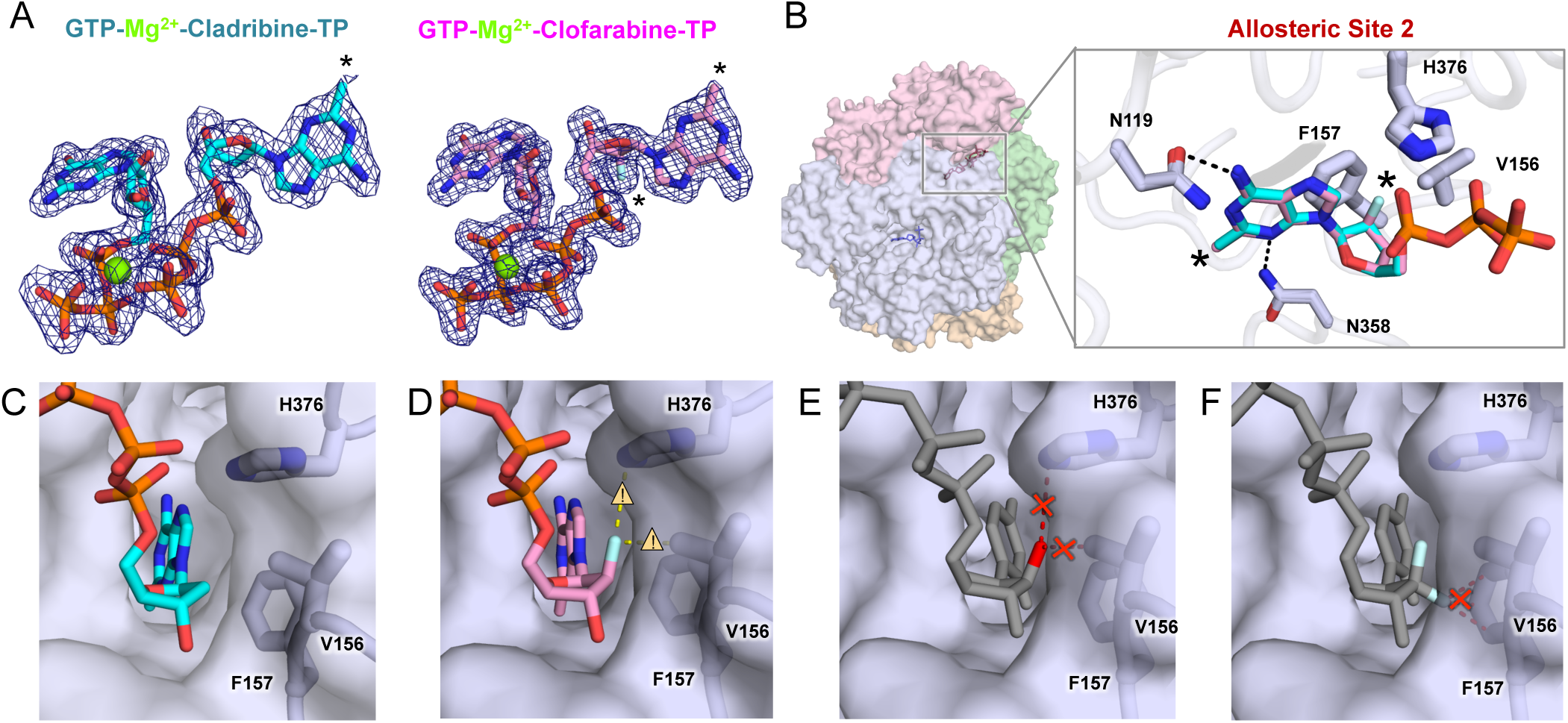
Structures of cladribine-TP and clofarabine-TP bound to Allo-site 2 of SAMHD1. (A) 2F_o_-F_c_ electron density (σ = 1.0) for GTP, cladribine-TP, and clofarabine-TP in the allosteric pocket of SAMHD1. Black asterisks indicate sites of modifications, and black dotted lines indicate hydrogen bonds. (B) Left: Transparent surface view of the SAMHD1 tetramer. Right: Overlay of cladribine-TP (cyan) and clofarabine-TP (pink) in Allo-site 2. Main chain of SAMHD1 is shown as tubes with selected residues and nucleotides represented as sticks. Residues important for gating the 2’-atom are highlighted in thicker sticks. Black asterisks indicate sites of modification. (C) The structure of cladribine-TP (cyan, sticks) in Allo-site 2 with V156, F157, and H376 shown as sticks under semitransparent surface of SAMHD1. (D) The structure of clofarabine-TP (pink, sticks) in Allo-site 2 with close contacts between the ((2’S)-2’-F) atom and V156, F157, and H376 highlighted with caution triangles and yellow dashed lines. (E) A model of cytarabine-TP in Allo-site 2 with potential steric clashes between the ((2’S)-2’-OH) group and V156 and H376 highlighted with a red cross and red dashed lines. (F) A model of gemcitabine-TP in Allo-site 2 with a potential steric clash between the ((2’R)-2’-F) atom and F157 highlighted with a red cross and red dashed lines.

While modest base modifications do not affect Allo-site 2 binding, modifications to the sugar at the 2’ position are restricted. We found that Allo-site 2 excludes all arabinose-based analogues with the ((2’S)-2’-OH) group, which is consistent with previous observations (31). The ribose-based ((2’R)-2’-F)-dCTP is also excluded from the site, the same as the NTP molecules reported before (19). As expected, Allo-site 2 does not tolerate a nucleotide with 2’,2’-difluorine modifications. The exclusion of nucleotides with these sugar modifications is likely due to SAMHD1 residues H376, V156, and F157 which form a tight pocket around the 2’ carbon of the dNTPs (Fig. 5B and 5C). While clofarabine-TP, which contains a ((2’S)-2’-F) modification in this position, is accommodated at this site, the structure shows that the fluorine atom may still cause some steric constraints (Fig. 5D). This is consistent with our oligomerization and activity assays, which showed reduced SAMHD1 activation by clofarabine-TP (Fig. 4). Modeling cytarabine-TP in Allo-site 2 pocket suggests steric clashes between the ((2’S)-2’-OH) group and residues H376 and V156 of SAMHD1 (Fig. 5E). On the other side of the sugar ring, residue F157 limits the accessibility of nucleotides with ((2’R)-2’-OH) or ((2’R)-2’-F) modifications, such as gemcitabine-TP (Fig. 5F). Cladribine-TP is the only deoxyribonucleotide analogue tested here, and most likely the lack of a 2’ sugar modification allows it to be the strongest allosteric activator of SAMHD1. Overall, Allo-site 2 in SAMHD1 was highly sensitive to modifications to the 2’ position of the sugar, and this moiety is a major binding determinant of nucleotide analogue binding to the allosteric site.

## Discussion

To improve current nucleotide analogue therapeutics, it is important to consider how these drugs interact with the cellular nucleotide metabolism pathways. As a negative regulator of the cellular nucleotide pool, SAMHD1 is an important cellular factor with the potential to influence nucleotide analogue efficacy. As is the case for cytarabine, SAMHD1 activity can convert the activated triphosphorylated form of the drug into its inactive precursor and consequently decrease the drug’s efficacy in patients (31). Conversely, if a nucleotide analogue drug is not a substrate of SAMHD1, then SAMHD1 depletes the cellular nucleotides competing for the same target sites, and thus increases the drug’s efficacy (33, 34). Therefore, understanding how nucleotide analogue drugs interact with SAMHD1, either as substrates or allosteric activators, has great potential for developing better cancer therapies. The study presented herein provides a comprehensive structural and biochemical framework for understanding how a wide range of nucleotide analogue drugs interact with SAMHD1. Characterizing these interactions will allow us to guide clinical practices by predicting how similar molecules might be affected by SAMHD1, and aid in the design of novel compounds with desirable properties.

Our biochemical and structural analysis of a panel of nucleotide analogues, with a variety of 2’ sugar modifications, reveals the detailed binding determinants for the catalytic site of SAMHD1. SAMHD1 selects substrates through indirect interactions between water molecules in the catalytic pocket and the base of the analogue drugs. The network of water molecules changes to adapt to variations found in the nucleotide analogues. Water-mediated interactions between the substrate and the enzyme allow for binding pocket plasticity and accommodation of different modifications. In addition, the catalytic pocket is accessible to the arabinose-like 2’ sugar modifications with the 2’S geometry. This provides a mechanistic understanding of the findings that arabinose-based nucleotides are substrates of SAMHD1 (31, 32). Interestingly, we were able to observe electron density for analogues with modified adenosine bases, which was previously unresolved in the available dATP-SAMHD1 structure (19). This indicates that the nucleotide analogue modifications may provide additional stabilizing interactions with the catalytic pocket of SAMHD1.

Our data also demonstrate that a nucleotide analogue can bind to the allosteric sites of SAMHD1 to activate the enzyme, depending on modifications to its sugar moiety (Fig. 5 and Fig. 6). We show that modest base modifications, such as a chlorine atom at C2 of adenine, are tolerated at the Allo-site 2. In contrast, Allo-site 2 does not allow arabinose-based or ribose-based nucleotides. Only deoxyribose-based nucleotides, such as cladribine-TP, can efficiently enter Allo-site 2 (Fig. 5 and 6). The results suggest that any modifications to the 2’ carbon of the sugar ring may affect the drug’s binding affinity to the allosteric pocket. These findings may help advance the understanding of the effect of current nucleotide analogue drugs and guide the design of non-hydrolysable analogues that can activate SAMHD1 for targeted depletion of cellular dNTP pools.

**Figure 6.**
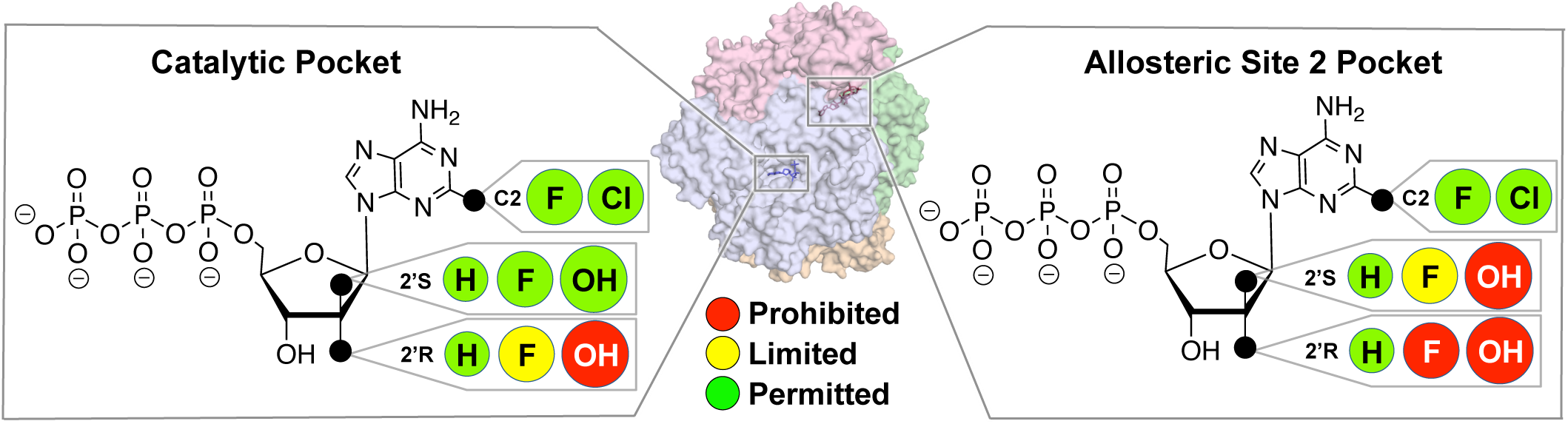
Summary of the effects of the 2’ sugar moiety on nucleotide analogue binding to the catalytic and allosteric sites of SAMHD1. Middle: Transparent surface view of SAMHD1 tetramer. Left: In the catalytic pocket of SAMHD1, while small substitutions such as fluorine atoms at the 2’R and 2’S positions of the sugar are permitted (green circle), their access to the 2’R position is limited (yellow circle). Larger modifications, such as hydroxyl groups, are permitted in the 2’S position, but not the 2’R position (red circle). Right: In Allo-site 2 of SAMHD1, hydroxyl groups are prohibited in both 2’R and 2’S positions of the sugar moiety. Fluorine atoms have limited access to the 2’S position, but they are prohibited from the 2’R position. Small base modifications such as fluorine or chlorine atoms are tolerated in both Allo-site 2 and the catalytic site.

Interestingly, we found that cytarabine-TP is particularly dependent on SAMHD1 cellular expression compared to other drugs tested in this study. Although many of the drugs tested were hydrolyzed by purified SAMHD1 in our biochemical assays, the effect of SAMHD1 expression on drug efficacy varied in cells. This may be due to different interactions between these drugs with other components of the nucleotide metabolism pathway in the cell. For example, even though clofarabine, fludarabine and cladrabine are similar in structure, these drugs have been reported to have varying degrees of interactions with cellular factors that affect their ability to be phosphorylated by dCK and to inhibit ribonucleotide reductase activity and DNA/RNA chain elongation (26). Moreover, nucleoside analogues may differentially influence activity and expression of cell cycle regulatory proteins such as cyclins and cyclin dependent kinases (39, 40) which in turn may interact with SAMHD1 and influence its dNTPase activity in leukemic cells (41). It remains possible that, although these drugs are hydrolyzed to a similar extent by SAMHD1 *in vitro*, their metabolism in the cell might be influenced by other cellular factors. In the case of cytarabine, our *in vivo* data strongly support that SAMHD1 is one of the main determinants of intracellular cytarabine-TP concentrations. On the other hand, we demonstrated that SAMHD1 strongly hydrolyzed cladribine-TP *in vitro* but had little influence on cladribine-TP concentrations and activity in AML cells. Combining biochemical, biophysical, and cellular studies offers a comprehensive approach to evaluate the response of nucleoside analogue drugs to SAMHD1 both at the mechanistic level and in application.

SAMHD1’s substrate promiscuity helps it function as a general sensor and regulator of nucleotide pools, but it also allows SAMHD1 to facilitate cancer cells to escape from nucleotide analogue treatments. Thus, it is important to consider how different modifications affect a nucleotide analogue drug’s access to the catalytic and allosteric sites of SAMHD1 (Fig. 6) when developing new therapies. Defining SAMHD1 interactions with nucleotide analogue drugs is critical for better predicting patient response to the current and future therapies.

## Materials and Methods

### Compounds for *in vitro* experiments

Cytarabine-TP, cladribine-TP, clofarabine-TP, fludarabine-TP, gemcitabine-TP, ((2’R)-2’-F)-dCTP and ((2’S)-2’-F)-dCTP were purchased from Jena Biosciences. GTP, dATP, and dCTP were purchased from Thermo Scientific.

### Protein expression and purification

N-terminal 6×His-tagged SAMHD1 constructs were expressed in *Escherichia coli* and purified using Ni-NTA affinity and size-exclusion chromatography as previously described (19).

### Analytical size exclusion chromatography

Purified samples of SAMHD1 (2 mg/ml, 50 µl) mixed with a final concentration of 500 µM GTP and 4 mM nucleotide analogue were applied to a Superdex 200 5/150 GL column (GE Healthcare) pre-equilibrated in 50 mM Tris-HCl, pH 8.0, 150 mM NaCl, 5 mM MgCl_2_ and 0.5 mM tris(2-carboxyethyl)phosphine (TCEP). The UV absorbance at 280 nm was measured as the protein sample eluted from the column.

### Analytical ultracentrifugation (AUC)

Sedimentation velocity experiments were performed with a Beckman XL-I analytical ultracentrifuge. Samples were prepared with protein concentration of 0.8-1.3 mg/ml in the buffer containing 50 mM Tris-HCl, pH 8.0, 150 mM NaCl, 5 mM MgCl_2_ and 0.5 mM TCEP and equilibrated with a final nucleotide concentration of 150 µM. AUC was performed at 35,000 revolutions per minute (r.p.m.) and 20°C with an An60-Ti rotor. The experimental parameters including sample partial specific volume, buffer density and viscosity were calculated with SEDNTERP (http://sednterp.unh.edu/). Velocity data were analyzed using the program SEDFIT (42).

### Crystallization and data collection

Purified SAMHD1 protein in buffer (50 mM Tris-HCl, pH 8.0, 150 mM NaCl, 5 mM MgCl_2_ and 0.5 mM TCEP) was mixed with 1 mM GTP and 10 mM analogue nucleotides (5 mM or 0.5 mM for gemcitabine-TP and vidarabine-TP, respectively) with or without 100 µM dATP (or 2.5 mM dATP for vidarabine-TP) and incubated at 4 °C for 15 minutes before crystallization. The small amount of dATP was included to ensure the formation of the SAMHD1 tetramer, as most nucleotide analogues do not bind Allo-site 2. All crystals were grown at 25 °C using the microbatch under-oil method by mixing 1 µL protein (5 mg/ml) with 1 µL crystallization buffer (100 mM SPG (Qiagen) buffer, pH 7.4, 25% PEG 1500). Crystals were cryoprotected by crystallization buffer supplemented with 25% (Vol/Vol) glycerol before frozen in liquid nitrogen. Diffraction data were collected at BNL beamline AMX and the Advanced Photon Source beamline 24-ID. The data statistics are summarized in Table 1.

### Structure determination and refinement

The structures were solved by molecular replacement using PHASER(43). We used the previously published SAMHD1 tetramer structure (PDB ID 4BZB), with the bound nucleotides removed, as the search model. The model was refined with iterative rounds of TLS and restrained refinement using *Refmac*5 (44) followed by rebuilding the model to the 2F_o_-F_c_ and the F_o_-F_c_ maps using Coot (45). Refinement statistics are summarized in Table 1. Coordinates and structure factors have been deposited in the Protein Data Bank, with accession codes listed in Table 1.

### Malachite green colorimetric assay

The enzymatic activity assay was adapted from (37). All assays were performed with purified catalytic domain of SAMHD1 (residues 113-626) at 25°C in a reaction buffer containing 50 mM Tris-HCl pH 8, 150 mM NaCl, 5 mM MgCl_2_, and 0.5 mM TCEP. Each 40 µL reaction, containing 10 µM pyrophosphatase, 0.5 µM SAMHD1, and 125 µM substrate or allosteric activator was quenched with 40 µL 20 mM EDTA after 15 minutes. Then, 20 µL Malachite Green reagent was added to the solution and developed for 15 minutes before the absorbance at 650 nm was measured.

### Compounds for *in vivo* experiments

Cytarabine (147-94-4), fludarabine (21679-14-1), clofarabine (123318-82-1) and cladribine (4291-63-8) were purchased from Tocris and gemcitabine (82092.00.00) from Accord Healthcare GmbH. All nucleotide standards, internal standards and dNs for the LC-MS/MS analysis were obtained from Sigma-Aldrich, Silantes or Alsachim (46). Cytarabine-^13^C_3_ (SC-217994) was purchased from Santa Cruz and used for LC-MS/MS analysis.

### Cells and cell culture

Human AML cell lines including THP-1 (DSMZ no. ACC16; FAB M6), OCI-AML2 (DSMZ No. ACC 99; FAB M4), OCI-AML3 (DSMZ No. ACC 582; FAB M4), Molm13 (DSMZ No. ACC 554; FAB M5a), PL-21 (DSMZ No. ACC 536; FAB M3), HL-60 (DSMZ No. ACC 3; FAB M2), MV4-11 (DSMZ No. ACC 102; FAB M5), SIG-M5 (DSMZ No. ACC 468; FAB M5a), ML2 (DSMZ No. ACC 15; FAB M4), NB4 (DSMZ No. ACC 207; FAB M3), KG1 (DSMZ No. ACC 14; FAB not indicated), MonoMac6 (DSMZ No. ACC 124; FAB M5), and HEL (DSMZ No. ACC 11; FAB M6) were obtained from DSMZ (Deutsche Sammlung von Mikroorganismen und Zellkulturen GmbH). THP-1 cells deficient for SAMHD1 (THP-1 KO) and corresponding control cells (THP-1 Control) were generated as previously described (47). All cell lines were cultured in IMDM (Biochrom) supplemented with 10 % FBS (SIG-M5 20% FBS), 4 mM L-Glutamine, 100 IU/ml penicillin, and 100 mg/ml streptomycin at 37°C in a humidified 5 % CO_2_ incubator. Cells were routinely tested for mycoplasma contamination (LT07-710, Lonza) and authenticated by short tandem repeat profiling, as reported (48).

### Cell viability assay

Viability of AML cell lines treated with various drug concentrations was determined by the 3-(4,5-dimethylthiazol-2-yl)-2,5-diphenyltetrazolium bromide (MTT) dye reduction assay after 96 hours of incubation as described previously (49). IC_50_ values were determined using CalcuSyn (Biosoft).

### Immunoblotting

Cells were lysed in Triton X-100 sample buffer and proteins separated by sodium dodecyl sulfate-polyacrylamide gel electrophoresis. Proteins were blotted onto a nitrocellulose membrane (Thermo Scientific). The membrane was incubated overnight at 4°C with primary antibodies used at the indicated dilutions: SAMHD1 (12586-1-AP, Proteintech, 1:1000), β-actin (3598R-100, BioVision via BioCat, 1:2000). Visualization and quantification was performed using fluorescently labeled secondary antibodies (926-32210 IRDye^®^ 800CW goat anti-mouse and 926-32211 IRDye^®^ 800CW goat anti-rabbit, LI-COR, 1:20000) and Odyssey LICOR.

### LC-MS/MS analysis

Cells (1 × 10^6^) were treated with 10 µM of the specific drug and incubated at 37 °C in a humidified 5% CO2 incubator for 6 h. Subsequently, cells were washed twice in 1 ml PBS, pelleted and stored at -20°C until measurement. The concentrations of dNTPs, ^13^C_3_-cytarabine-TP, fludarabine-TP, clofarabine-TP, cladribine-TP and gemcitabine-TP, in the samples were measured by liquid chromatography-electrospray ionization-tandem mass spectrometry as previously described (46). Briefly, the analytes were extracted by protein precipitation with methanol. An anion exchange HPLC column (BioBasic AX, 150 × 2.1 mm, Thermo) was used for the chromatographic separation and a 5500 QTrap (Sciex) was used as analyzer, operating as triple quadrupole in positive multiple reaction monitoring (MRM) mode. The analysis of the dNTP was performed as previously described (46). Additionally, ^13^C_3_-cytarabine-TP, fludarabine-TP, clofarabine-TP, cladribine-TP and gemcitabine-TP were quantified using cytidine-^13^C_9_ -^15^N_3_-5’-triphosphate as internal standard (IS). The precursor-to-product ion transitions used as quantifiers were: m/z 487.0 → 115.1 for ^13^C_3_-cytarabine-TP, m/z 525.7 → 154.1 for fludarabine-TP, m/z 543.7 → 134.0 for clofarabine-TP, m/z 526.0 → 170.0 for cladribine-TP and m/z 504.0 → 326.0 for gemcitabine-TP. Due to the lack of commercially available standards, relative quantification was performed by comparing the peak area ratios (analyte/IS) of the differently treated samples.

### Production of virus-like particles (VLPs)

VLPs, carrying either Vpx or Vpr from SIVmac251, were produced by co-transfection of 293T cells with pSIV3+ gag pol expression plasmids and a plasmid encoding VSV-G. The SIVmac251-based gag-pol expression constructs pSIV3+R-(Vpr-deficient) and pSIV3+X-(Vpx-deficient) were previously reported (50). The SAMHD1 degradation capacity of Vpx-VLPs was determined in THP-1 cells 24 h post transduction by intracellular SAMHD1 staining. AML cell lines were spinoculated with VSV-G pseudotyped VLPs carrying either Vpx or Vpr. Expression of SAMHD1 was monitored by Western blotting.

### Statistical Information

The average standard errors and standard deviations were calculated from multiple separate experiments as indicated in each figure legends and the results are shown in each graph.

### Data deposition

The atomic coordinates and structure factors have been deposited in the Protein Data Bank, www.pdb.org (PDB ID codes XXX, XXX, XXX, XXX, XXX).

## Acknowledgments

We thank J. Wang for technical assistance and discussions. We also thank the staff at the Advanced Photon Source beamlines 24-ID. This work was supported in part by NIH grants AI102778 (Y.X) and AI120845 (X.J.), by the Yale Cancer Center Pilot Grant (Y.X.), by Hilfe für krebskranke Kinder Frankfurt e.V. and the Frankfurter Stiftung für krebskranke Kinder (J.C and C.S), and by the Deutsche Forschungsgemeinschaft (KE 742/4-1) (O.T.K). O.B. was supported by the predoctoral program in Cellular and Molecular Biology NIH T32 GM007223 and by the National Science Foundation Graduate Research Fellowship DGE1122492. K.M.K. was supported by the predoctoral program in Biophysics NIH T32 GM008283. G.G. was supported by the Hessischen Landesoffensive zur Entwicklung wissenschaftlicher und ökonomischer Exzellenz (LOEWE-Center) Translational Medicine and Pharmacology.

---

**Supplementary Figure S1.**
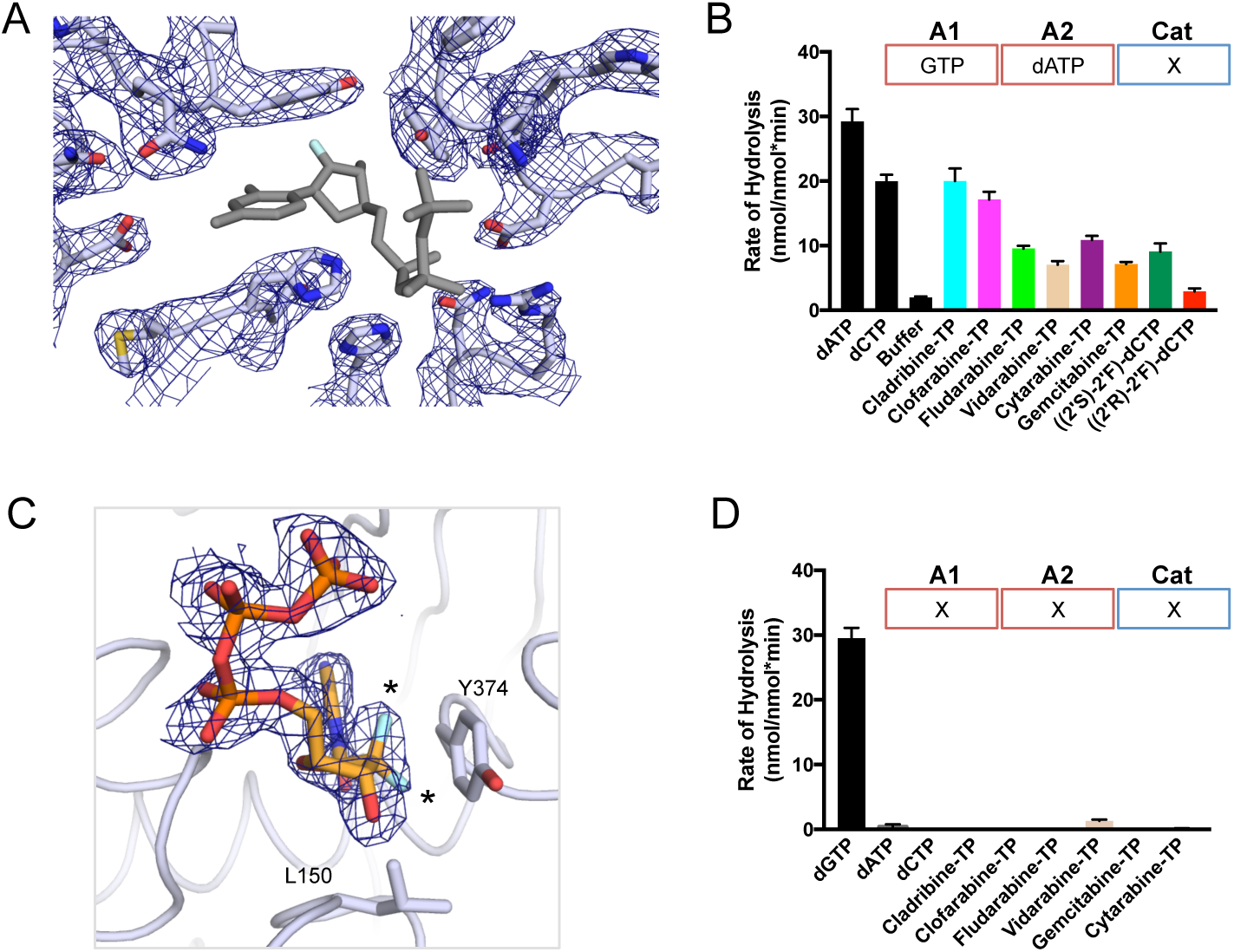
The catalytic and allosteric pockets exhibit different preferences for nucleotides. (A) 2F_o_-F_c_ electron density (σ = 1.0) in the catalytic pocket of SAMHD1. No electron density is observed for ((2’R)-2’-F)-dCTP (modeled as grey sticks). (B) dNTPase assay measuring triphosphates produced by pre-assembled SAMHD1 tetramer in the presence of 125 uM dCTP, dATP, nucleotide analogues, or buffer alone. Error bars represent SEM. (C) 2F_o_-F_c_ electron density (σ = 1.0) for gemcitabine-TP in the catalytic pocket with labeled residues shown in sticks and black stars indicate the sites of modifications. (D) dNTPase assay measuring triphosphates produced by SAMHD1 mixed with 125 uM dGTP, dCTP, dATP, nucleotide analogues, or buffer alone. Error bars represent SEM of three independent experiments.

**Supplementary Figure S2.**
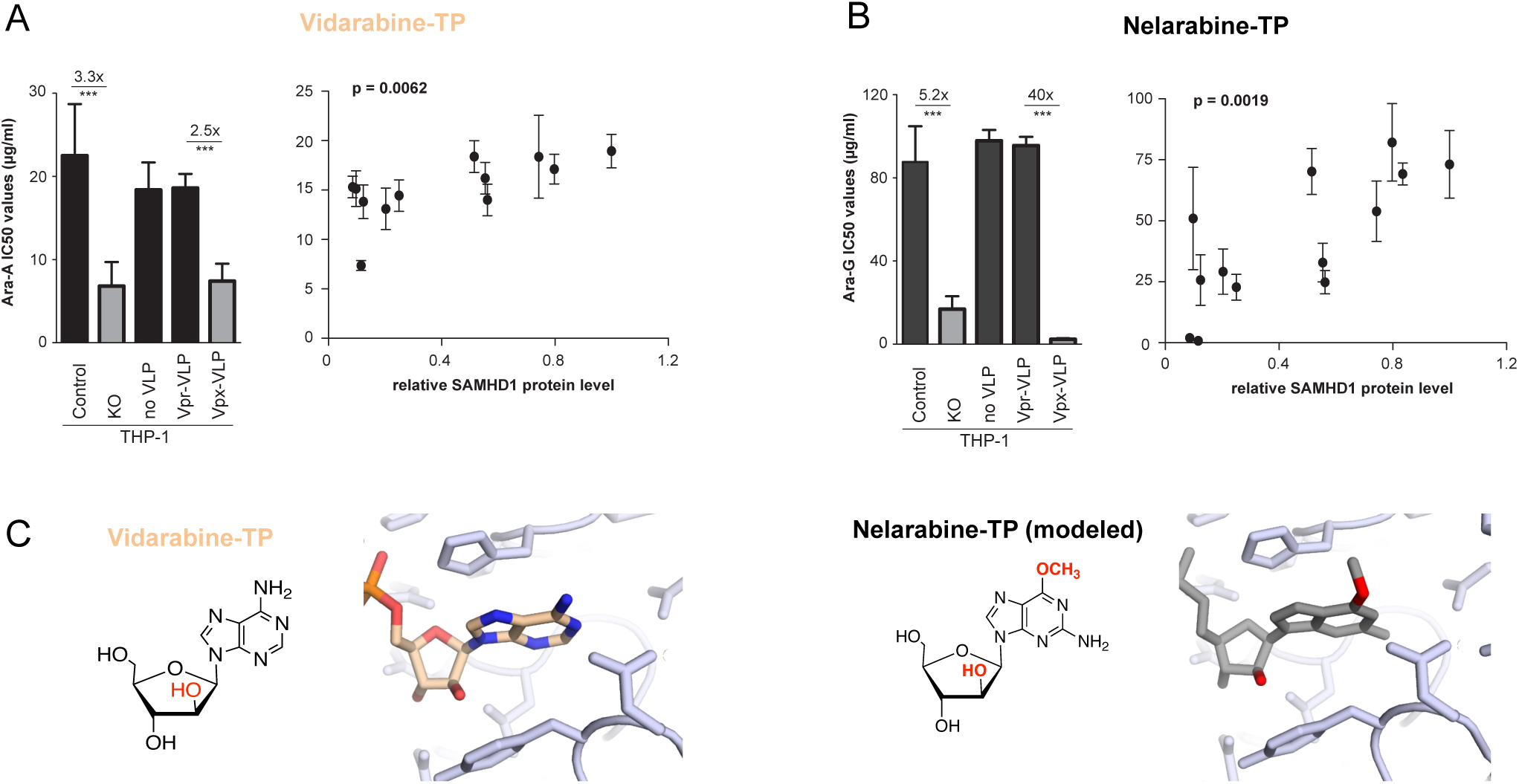
SAMHD1 affects vidarabine-TP and nelarabine-TP cytotoxicity *in vivo*. Left: (A) Vidarabine-TP and (B) nelarabine-TP IC_50_ values in THP-1 KO cells or THP-1 cells transduced with VLPs carrying either lentiviral Vpr (Vpr-VLPs, control) or Vpx proteins (Vpx-VLPs) shown in bars representing mean ± SEM of three independent experiments. Numbers indicate the factor of decrease of the IC_50_ values in the absence of SAMHD1. Right: Correlations of vidarabine-TP (A) and nelarabine-TP (B) concentrations inhibiting 50% of cell viability (IC_50_) and relative protein expression levels of SAMHD1 in 13 AML cell lines. Relative expression levels (ratios of a SAMHD1/β-actin) are shown as arbitrary units (a.u.). Ratio of SAMHD1/β-actin for THP-1 was set to 1 and ratios of other cell lines set relative to it. Closed circles represent mean ± SEM of three independent experiments. (C) Left: Vidarabine chemical structure and crystal structure of vidarabine-TP bound to the catalytic pocket of SAMHD1. SAMHD1 backbone is shown as coils with side chains and vidarabine-TP shown as sticks. Right: Nelarabine chemical structure and nelarabine-TP modeled into the catalytic pocket of SAMHD1.

Author contributions
K.M.K., O.B., C.S., D.T., N.F., J.C., O.T.K., and Y.X. designed research; K.M.K., O.B., C.S., D.T., and V.S. performed research; K.M.K., O.B., C.S., D.T., F.T., K.R., N.F., G.G., V.B., X.J., O.T.K., J.C., and Y.X. analyzed data; and K.M.K., O.B., and Y.X. wrote the paper with contributions from O.T.K. and J.C.

## Notes

The authors declare no conflict of interest.

